# Exploring a local genetic interaction network using evolutionary replay experiments

**DOI:** 10.1101/2021.01.27.428500

**Authors:** Ryan C. Vignogna, Sean W. Buskirk, Gregory I. Lang

## Abstract

Understanding how genes interact is a central challenge in biology. Experimental evolution provides a useful, but underutilized, tool for identifying genetic interactions, particularly those that involve non-loss-of-function mutations or mutations in essential genes. We previously identified a strong positive genetic interaction between specific mutations in *KEL1* (P344T) and *HSL7* (A695fs) that arose in an experimentally-evolved *Saccharomyces cerevisiae* population. Because this genetic interaction is not phenocopied by gene deletion, it was previously unknown. Using “evolutionary replay” experiments we identified additional mutations that have positive genetic interactions with the *kel1*-P344T mutation. We replayed the evolution of this population 672 times from six timepoints. We identified 30 populations where the *kel1*-P344T mutation reached high frequency. We performed whole-genome sequencing on these populations to identify genes in which mutations arose specifically in the *kel1*-P344T background. We reconstructed mutations in the ancestral and *kel1*-P344T backgrounds to validate positive genetic interactions. We identify several genetic interactors with *KEL1*, we validate these interactions by reconstruction experiments, and we show these interactions are not recapitulated by loss-of-function mutations. Our results demonstrate the power of experimental evolution to identify genetic interactions that are positive, allele specific, and not readily detected by other methods, and sheds light on a previously under-explored region of the yeast genetic interaction network.

## Introduction

Genetic interactions drive evolutionary change (Breen et al. 2012), define functional relationships between genes (Costanzo et al. 2010; Costanzo et al. 2016), and contribute to disease, behaviors, and other complex traits (Manolio et al. 2009; Zuk et al. 2012). Despite the importance of genetic interactions, we lack a broad understanding of their prevalence and their properties. Most of our understanding of genetic interactions comes from systematic screens of double deletion mutants (Tong et al. 2004; Baryshnikova et al. 2010; Babu et al. 2014; Costanzo et al. 2016; Kuzmin et al. 2018). Double deletion analyses and alternative approaches such as RNAi (Bassik et al. 2013) or CRISPR/Cas9 library screens (Han et al. 2017; Shen et al. 2017) and transposon mutagenesis (Horton and Kumar 2015) focus almost exclusively on loss-of-function mutations, which are rarely beneficial and account for only a small fraction of standing natural variation (Bergström et al. 2014; Saleheen et al. 2017). Conversely, methods that assess the phenotypic effects of natural variants, such as genome-wide association studies and quantitative trait locus (QTL) mapping experiments, are underpowered to detect genetic interactions (Marchini et al. 2005; Jiang et al. 2010; Zuk et al. 2012; Fang et al. 2019). Experimental evolution can bridge the gap between systematic deletion analyses and QTL mapping by identifying genetic interactions between mutations that are representative of natural genetic variation including non-loss-of-function mutations and mutations in essential genes.

Genetic interactions are thought to be prevalent among fixed mutations in experimental evolution (Chou et al. 2011; Khan et al. 2011; Buskirk et al. 2017). Previous experiments have identified genetic interactions statistically, most often as pairs of genes in which mutations co-occur more (or less) often than expected by chance across replicate populations (Kvitek and Sherlock 2011; Tenaillon et al. 2012; Good et al. 2017; Fisher et al. 2019). Most interactions identified in this way are negative genetic interactions, where the combined effect of mutations is less beneficial than expected given the individual mutations. In contrast, relatively few positive genetic interactions, in which the combined effect of mutations is more beneficial than expected, have been identified and characterized (Blount et al. 2008; Quan et al. 2012; Buskirk et al. 2017; Fisher et al. 2019). It is possible that beneficial positive genetic interactions are less common or, instead, they may be more difficult to detect by existing methods (Kryazhimskiy et al. 2009; Khan et al. 2011; Kryazhimskiy et al. 2014).

Previously, we identified a positive genetic interaction between two mutations that arose several hundred generations apart during the course of a yeast laboratory evolution experiment (Buskirk et al. 2017). One of the first mutations to arise was a missense mutation in *KEL1* (P344T) that had little effect on fitness by itself. Later, a frameshift mutation in *HSL7* (A695fs) arose that had a much larger fitness effect on the *kel1*-P344T background than it would have had on the ancestral background. These two genes, *KEL1* and *HSL7* were not previously known to interact genetically and this interaction may have been inaccessible to methods that systematically survey pairwise loss-of-function mutations.

Here we use evolutionary-replay experiments to test the hypothesis that the *kel1*-P344T mutation “opened the door” to mutations that would otherwise be less favored. By restarting the evolution of this population hundreds of times we allow selection to enrich for mutations that, like the *hsl7*-A695fs mutation, interact positively with the *kel1*-P344T mutation. By initiating replays from timepoints before and after the *kel1-*P344T mutation arose we identify genes that fix preferentially in the *kel1*-P344T background. We identify both known and previously unknown *KEL1* genetic interactors and validate putative genetic interactions by genetic reconstruction. We show interactions identified in this way are positive, beneficial, and unlikely to be uncovered by systematic analysis of double deletions or knockdowns.

## RESULTS

### Evolutionary replay experiments reveal the range and likelihood of alternative outcomes

Previously, we identified a strong positive genetic interaction that arose in a 1,000-generation laboratory-evolved yeast population (Lang et al. 2013). In this population (BYS2-E01 from Lang et al. 2011) a low frequency lineage persisted for hundreds of generations and acquired a *KEL1* mutation (P344T) by Generation 335 (Figure 1A). We showed that the *kel1*-P344T mutation is nearly neutral in the ancestral background and the success of this low-frequency lineage (referred to as our focal lineage) was due to a mutation in *HSL7* (A695fs) that arose before Generation 600 and had a strong positive genetic interaction with *kel1*-P344T (Figure 1B) (Buskirk et al. 2017). This positive genetic interaction allowed our focal lineage to outcompete an ascendant *ste12* lineage present in the population.

**Figure 1.**
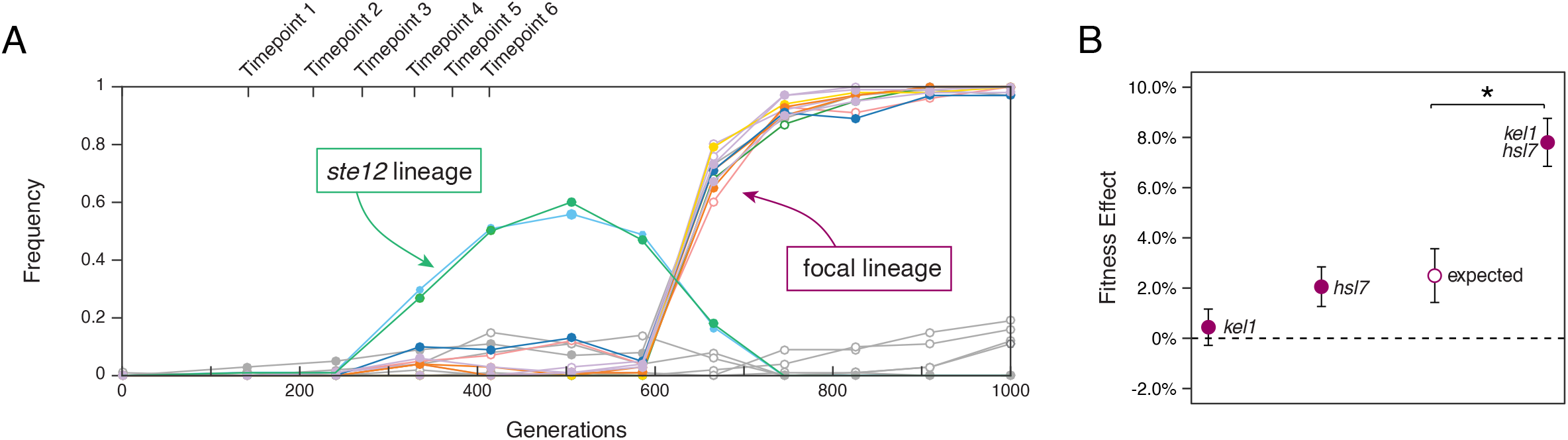
Dynamics of adaptation of population BYS2-E01. (A) Frequency of mutations present in the original BYS2-E01 population as determined by whole genome sequencing (Lang et al. 2013). A lineage containing a beneficial *ste12*-Q151fs mutation was outcompeted when an *hsl7*-A695fs mutation arose on the low-frequency focal lineage. Replay experiments were initiated at the timepoints indicated above (Generations 140, 210, 270, 335, 375, and 415). The *kel1*-P344T mutation was first detected by whole genome, whole population sequencing at Timepoint 4. (B) Average fitness effects and standard deviations of mutations that contribute to the fitness of the focal lineage (Buskirk et al. 2017). Expected double mutant fitness (open circle) was calculated by summing individual fitness effects and propagating uncertainty. Asterisk (*) indicates p < 0.001 (Welch’s modified t-test).

To investigate the role of the *kel1*-P344T mutation on the fate of our focal linage we resurrected the population in which it arose from timepoints before and after the *kel1*-P344T mutation was detected: Generations 140, 210, and 270 and Generations 335, 375, and 415, respectively (Figure 1A). For each timepoint we restarted 48 replicate replay populations and evolved them for 630 generations under the same conditions as the original experiment, with daily 1:2^10^ dilutions into 128 μL of rich glucose medium (see Methods).

Each week (every 70 generations), we followed the fate of our focal lineage in all 288 replay populations using a two-step process: first by screening phenotypically for loss of the *ste12* lineage using a fluorescent reporter of Ste12 activity, which is present in this strain background (Lang et al. 2011), then by genotyping for a mutation present only in our focal lineage. Since the *ste12*-Q151fs and *kel1*-P344T mutations arose on independent backgrounds, only in populations where the *ste12* lineage goes extinct could our focal lineage have fixed. We find that the *ste12* lineage “won” (either fixed or was at a frequency above 0.5 at Generation 630) in most populations from both the pre-*kel1* (118 of 144) and post-*kel1* (127 of 144) timepoints.

For those populations where the *ste12* lineage “lost” (either went extinct or was at a frequency below 0.5 at Generation 630), we determined if our focal lineage had won. For this, we took advantage of a *SspI* restriction site that was introduced by the *iqg1*-S571N mutation that arose in our focal lineage around Generation 140, prior to the *kel1*-P344T mutation (note that, although the *kel1*-P344T mutation is only present in replay experiments initiated at Generation 335 and beyond, the *iqg1*-S571N mutation identifies our focal lineage in all six replay experiments and has no detectable effect on fitness (Buskirk et al. 2017)) (Figure S1). In only one of the pre-*kel1* replay populations did our focal lineage win. However, in the post-*kel1* replay populations, our focal (*iqg1*-S571N, *kel1*-P344T) lineage won in 4 of the 17 populations where the *ste12* lineage lost (Figure 2, Table S1).

**Figure 2.**
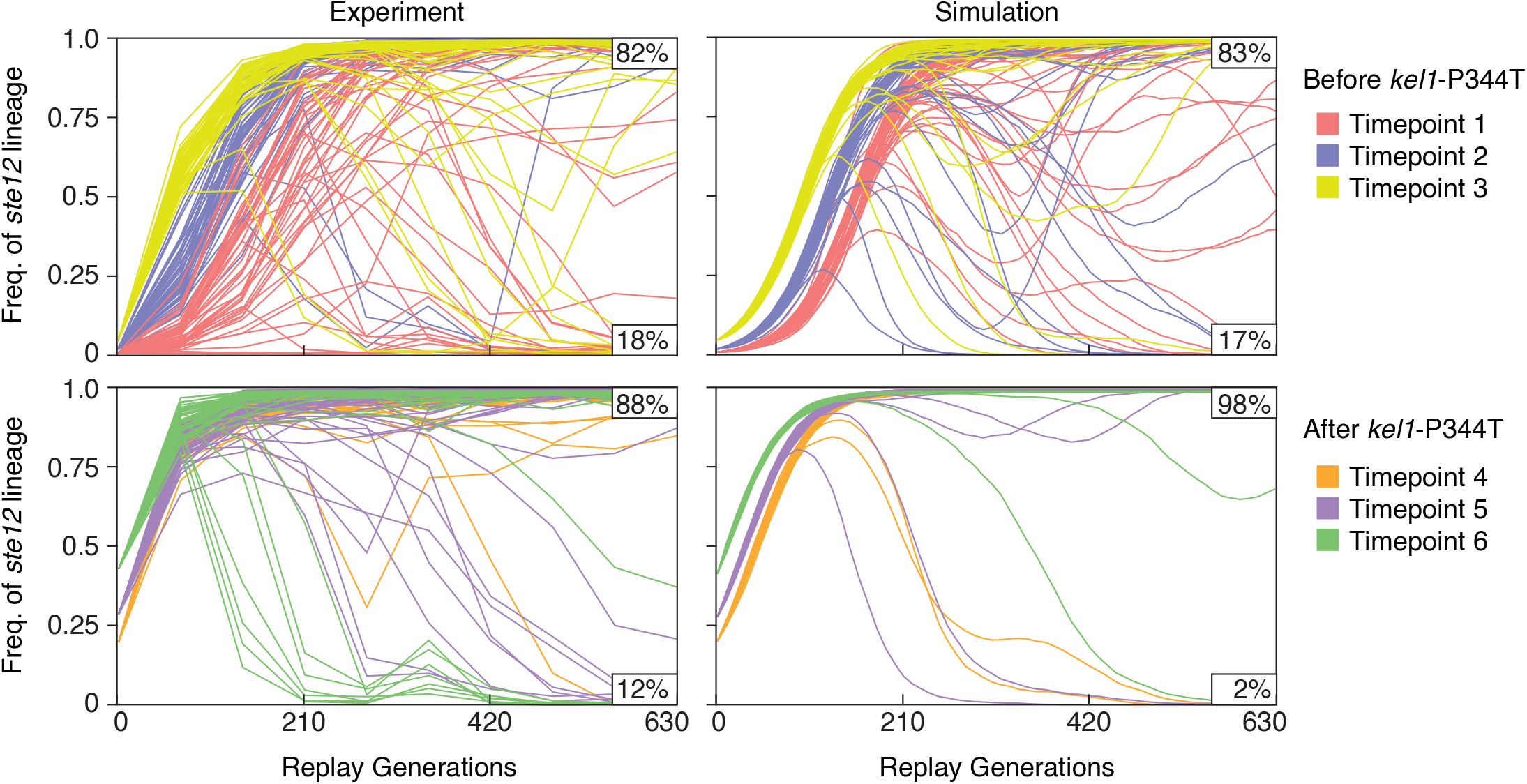
Fate of *ste12* lineages in the replay populations and simulated populations. Frequency of sterile cells in the replay populations over the course of the replay experiments as measured by flow cytometry (left) and frequency of the *ste12* lineage over time in simulated populations (right). Each line represents one population. Percentage of populations where the *ste12* lineage wins (top value) or loses (bottom value) are shown for each graph. For the simulations we show the first 48 simulated trajectories as an unbiased representation. Initial frequencies of *ste12* lineage in experimental populations were plotted from values reported in Lang et al. 2011.

### Chance alone is insufficient to explain observed lineage dynamics

We sought to determine whether the appearance of the neutral *kel1*-P344T mutation in our low-frequency focal lineage affected the likelihood that the dominant *ste12* lineage won or lost in the replay populations. Direct comparison between timepoints is not possible because each subsequent set of replay experiments starts with a higher initial frequency of the *ste12* lineage (Figure 1A). To address this, we simulated the evolutionary replay experiments using a previously-described computational model of our experiment (Frenkel et al. 2014).

For each of the initial frequencies of the *ste12* lineage, we ran 10,000 forward evolution simulations using known values for the starting frequency of the *ste12* lineage (Lang et al. 2013), the fitness of the *ste12* lineage, and empirically-derived values for the beneficial mutation rate and the distribution of fitness effects in our experiment (see Methods). In the simulations we find that the fraction of replay populations where the *ste12* lineage wins is greater for each subsequent replay timepoint (Table S1). This is expected since the *ste12* lineage starts at a higher frequency in each successive replay timepoint from <0.01 at Generation 140 to 0.42 at Generation 415.

In our experiment, like in the simulations, the *ste12* lineage wins the fewest times for replays initiated at the first timepoint when the *ste12* lineage starts at a frequency of <0.01. Apart from the first timepoint, however, the *ste12* lineage wins the fewest times in the final two replay timepoints (Timepoints 5 and 6) where the *ste12* lineage started at its highest frequencies (0.28 and 0.42) and where in the simulations the *ste12* lineage won most often. The key difference between the simulation and the experiment is that the simulation assumes that the distribution of fitness effects of new beneficial mutations is fixed. In reality, this distribution will be impacted by the presence of other mutations in the population, such as the low-frequency *kel1*-P344T mutation that arose around Generation 335. Grouping the first three (pre-*kel1*) timepoints we find the likelihood of the *ste12* lineage winning in the replay experiments does not deviate from the null model given by the simulations (*X*^2^ = 0.05, p = 0.82). However, grouping the last three (post-*kel1*) timepoints we find that the *ste12* lineage loses more often than expected by chance (*X*^2^ = 63, p < 0.001) (Figure 2, Table S1).

Our hypothesis for why the *ste12* lineage lost more often than expected in replay experiments initiated after Generation 335 is that the presence of the *kel1*-P344T mutation opens paths to higher-fitness mutations (e.g., the fitness effect of the *hsl7*-A695fs mutation is greater in the background of the *kel1*-P344T mutation). It would follow then that the deviations from the null expectation are due to populations where our focal lineage won. However, if we exclude from our analysis those populations where our focal lineage won, we find that the *ste12* lineage still loses more often than expected by chance in the later three timepoints (*X*^2^ = 34, p < 0.001). Therefore, accounting for the *kel1*-P344T mutation can explain some, but not all, of the deviations between the simulated and actual replay experiments. This suggest that additional low-frequency lineages in the population influence the dynamics of adaptation (see Discussion).

### Recurrent targets of selection are putative *KEL1*-interacting genes

Genes that acquire mutations more often than expected by chance only in populations where our focal lineage won are candidate genetic interactors with *kel1*-P344T. Because our focal lineage won in only 4 of the 144 populations in the initial replay experiments, we initiated an additional 384 replay populations from Timepoint 5 (in which our focal lineage won 3 out of 48 times) and evolved them for 350 generations. We used the same two-step method for identifying populations where our focal lineage won: screening for loss of the *ste12* lineage followed by PCR/*SspI* digest to detect the presence of the *iqg1*-S571N mutation. The *ste12* lineage lost in 45 of the 384 second-round replay populations and our focal lineage won in 26 of those 45.

We sequenced one clone from each population in which our focal lineage had won: 5 populations from the first round of replay experiments and 26 from the second round. Additionally, we sequenced 19 randomly selected control populations that saw either the *ste12* lineage or another non-*ste12*, non-focal lineage win (Supplemental Dataset 1). We identified 526 *de novo* mutations that arose during the replay experiments; an average of ∼13 mutations per clone in the first-round replays and an average of ∼9 per clone in the second-round replays. Autodiploidization events, a common outcome in our experimental system (Fisher et al. 2018), were detected in 17 out of the 50 sequenced clones (Supplemental Dataset 1).

We classified mutations as putative *kel1*-P344T interacting mutations if they occurred in genes previously known to interact with *KEL1* and/or in genes with multiple unique mutations in independent replay populations. A total of 22 genes fulfill the first criterion, and four genes fulfill the second. One gene, *HSL1*, fulfills both criteria (Supplemental Dataset 1).

### Reconstruction experiments validate positive genetic interactions

We reconstructed putative *kel1*-interacting mutations in both the ancestral and *kel1*-P344T backgrounds. We included mutations in known *KEL1* interacting genes, recurrent targets of selection, and/or essential genes, as well as a gene mutated in both *kel1* and non-*kel1* lineages as a negative control. Each mutation was assayed for fitness (alone and in combination with the *kel1*-P344T mutation) against a fluorescently-labeled ancestral reference strain (Supplemental Dataset 2).

*HSL1* is a both a known *KEL1* interactor (Sharifpoor et al. 2012; Kryazhimskiy et al. 2014; Kuzmin et al. 2018) and a recurrent target of selection, mutated in three independent replay populations. By reconstructing the *hsl1*-A262P mutation on the *KEL1* background we find that it has a 3.50 ± 0.3% (α = 0.05) fitness effect. On the *kel*1-P344T background, however, it has a 6.53 ± 0.2% (α = 0.05) fitness effect, exceeding the additive expectation (p < 0.001 Welch’s modified t-test, Figure 3A). A mutation (D233fs) in another known *KEL1* interactor, *PSY2*, is also more beneficial in the presence of *kel1*-P344T than in its absence (p < 0.001 Welch’s). Interestingly, in this second interaction, both the *kel1*-P344T and *pys2-*D233fs mutations are neutral in isolation yet produce a 3.48 ± 0.3% (α = 0.05) benefit when combined.

**Figure 3.**
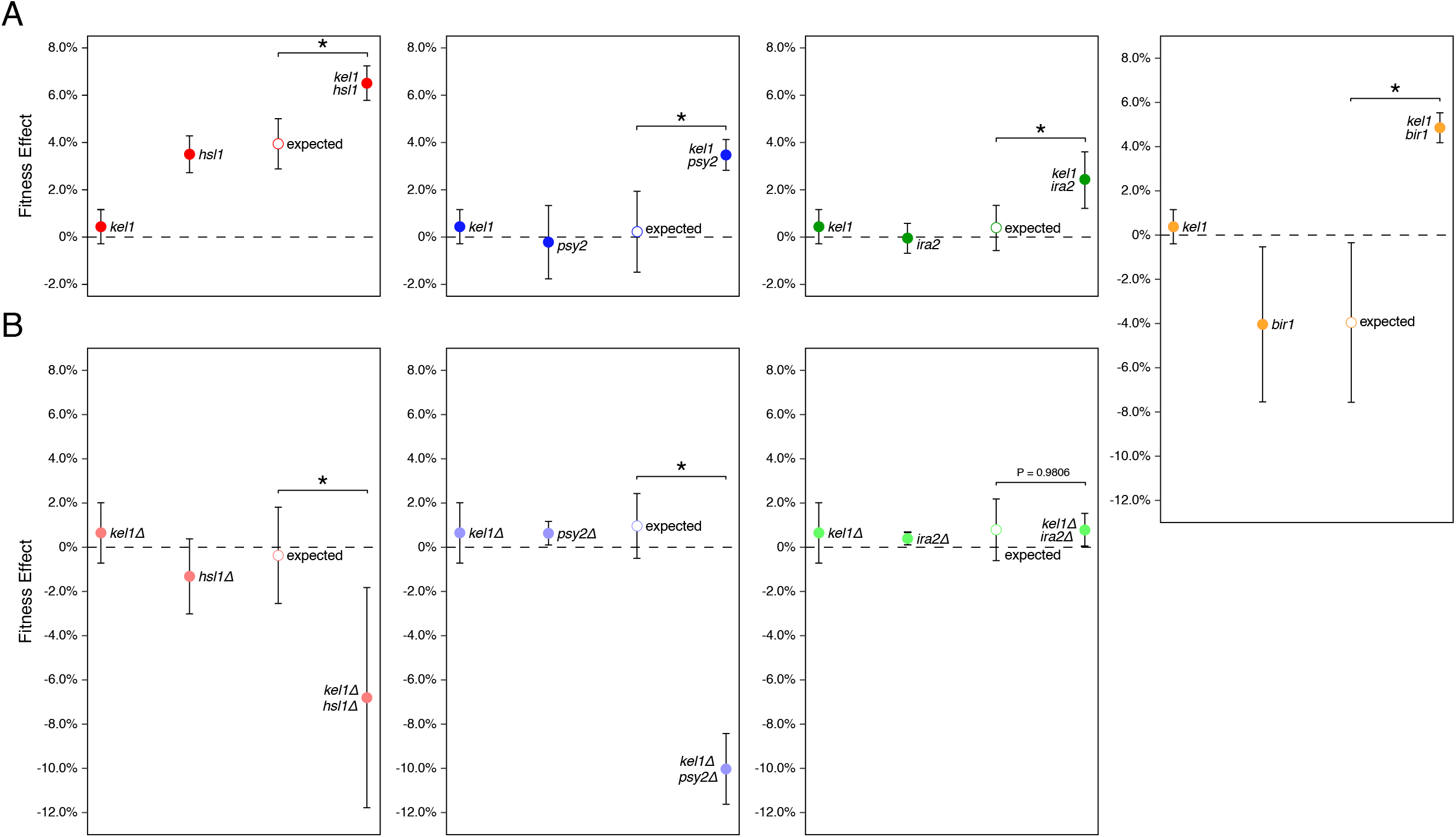
Positive genetic interactions between evolved mutations. Average fitness effect and standard deviation of (A) evolved mutations when reconstructed in the ancestral background and (B) gene deletions in the ancestral background. Observed fitness effects were determined by competitive fitness assays against a fluorescently-labeled version of the ancestor. Expected double mutant fitness (open circles) were calculated by summing individual fitness effects and propagating uncertainty. Asterisk (*) indicates p < 0.001 (Welch’s modified t-test).

In addition to validating two known *KEL1* interactors, we identify two genes that interact genetically with *kel1*-P344T but were not previously recognized as *KEL1* interactors (Figure 3A). A nonsense mutation in *IRA2* (R1575^*^) has a fitness cost of −0.47 ± 0.2% (α = 0.05) in the ancestral background but has a 2.44 ± 0.4% (α = 0.05) fitness advantage when combined with the *kel1*-P344T mutation (p < 0.001 Welch’s). A *BIR1* mutation (P685T) has a substantial fitness cost of −4.51 ± 0.7% (α = 0.05) in the ancestral background but confers a fitness benefit of 4.26 ± 0.3% (α = 0.05) when combined with the *kel1*-P344T mutation (p < 0.001 Welch’s). Neither the fitness effect of a control mutation in *ECM21* (P799fs) nor a mutation in *YLR001C* (R427fs), a recurrently mutated gene in the replay experiments, are influenced by the *kel1*-P344T mutation (p = 0.2060 and p = 0.0347 respectively, Welch’s, Figure S2). Taken together, these results highlight the ability of our experimental system to identify known and previously unknown genetic interactions.

### Genetic interactions revealed through experimental evolution are positive and allele specific

Experimental evolution samples all types of spontaneous mutations including loss, gain, or attenuation of function. As such, interactions identified in this way may be allele specific. We tested for allele specificity by determining whether the interactions we identified could be phenocopied by gene deletion. For each *KEL1* interactor, we generated full-gene deletions in the ancestral background as well as in an isogenic *KEL1* gene deletion (*kel1*Δ) mutant background (Figure 3B). Like the *kel1*-P344T mutation, the *kel1*Δ mutation by itself is neutral.

*HSL1* and *PSY2* have previously been identified as *KEL1* genetic interactors using double-deletion mutants in yeast (Costanzo et al. 2016; Kuzmin et al. 2018). We also observe genetic interactions between *kel1Δ* and *hsl1Δ* as well as between *kel1Δ* and *psy2Δ* (both p < 0.001, Welch’s, Figure 3B). Importantly, however, the direction of the interactions differs between the gene deletions (which decrease fitness) and the evolved mutations (which increase fitness). Therefore, although *KEL1/HSL1* and *KEL1/PSY2* genetic interactions can be identified by gene deletion, the effects of these interactions on fitness are allele specific.

Although we observe a positive genetic interaction between evolved mutations in *IRA2* and *KEL1* we do not detect a genetic interaction between *kel1Δ* and *ira2Δ* (p = 0.9806, Welch’s, Figure 3B). A *bir1Δ* mutation was not constructed because it is an essential gene (Li et al. 2000; Widlund et al. 2006). Together, these results demonstrate the power of experimental evolution to identify genetic interactions that are not easily detectable by current methods.

## DISCUSSION

Genetic interactions impose constraints on the paths that evolution can take, and thus they determine the extent to which the evolutionary future of a population is dependent on its past. Retrospective analysis of the mutations that arose during laboratory experimental evolution has found evidence for both negative (Chou et al. 2011; Khan et al. 2011; Tenaillon et al. 2012; Kryazhimskiy et al. 2014) and positive genetic interactions (Blount et al. 2008; Quandt et al. 2015; Buskirk et al. 2017; Fisher et al. 2019). Here, we reverse the standard paradigm. Rather than identifying genetic interactions among mutations that arose during experimental evolution, we use experimental evolution as a tool to recover mutations that interact with a specific mutation of interest.

Experimental evolution provides an opportunity to identify genetic interactions that are potentially overlooked by current methods. Unlike suppressor screens, which require a selectable phenotype, continuous selective pressure exerted over hundreds of generations efficiently enriches for genotypes that contain positive genetic interactions whose effects may be subtle. Mutations arise spontaneously and cover all types of mutations including non-loss-of-function mutations and mutations in essential genes, making for a much larger and diverse search space, and one that is likely more reflective of natural genetic variation.

We previously detected a positive genetic interaction between mutations in *KEL1* and *HSL7* that arose during a laboratory evolution experiment (Lang et al. 2013; Buskirk et al. 2017). Though their protein products both localize to sites of polarized growth, these two genes were not previously known to interact physically or genetically. Two high-throughput screens have identified possible interactions between *KEL1* and other cell cycle genes, yet the local genetic interaction network linking the cell polarity gene *KEL1* and the cell cycle has not been fully explored (Ho et al. 2002; Ubersax et al. 2003).

Here we use evolutionary replay experiments to identify other genetic interactors that, like *hsl7*-A695fs, have a significantly higher fitness effect on the *kel1*-P344T background. The population used here, BYS2-E01 from Lang et al. 2011, is ideal for this purpose for two reasons: (1) the nearly neutral *kel1*-P344T mutation arose and persisted at low frequency for several hundred generations prior to the *hsl7*-A695fs mutation, and (2) the presence of the beneficial *ste12*-Q151fs mutation and the restriction site created by the *iqg1*-S571N mutation allows individual lineages of interest to be tracked quickly and easily without sequencing.

We followed the fate of *ste12*-Q151fs across 48 replicate replay experiments initiated at Generations 140, 210, 270, 335, 375, and 415. All else being equal, the *ste12* lineage should win more often in populations initiated at later timepoints since this lineage starts at higher frequencies in each subsequent timepoint. We verified this null result by computational simulations that capture the details of our experiment (Figure 2). We predicted that after Timepoint 4 the evolutionary potential of our focal lineage would increase due to new beneficial mutations made accessible by the *kel1*-P344T mutation. Therefore, we expected to see a reduction in the fraction of replay populations in which *ste12* lineage won in our experiment relative to the simulations. Comparing our replay data to the simulations, we find that the *ste12* lineage loses more often than expected in post-*kel1* timepoints of the replay experiments. Though this difference is significant, the magnitude of the effect is not strong. This could indicate that the additional beneficial mutations available to the *kel1*-P344T-containing cells does not appreciably alter the distribution of fitness effects. Alternatively, the *kel1*-P344T mutation may “close” as many paths to higher fitness as it “opens,” resulting in a shift in the identity of beneficial mutations but not the distribution of fitness effects.

The deviations between experiment and simulation cannot be due only to the appearance of the *kel1*-P344T mutation. If we exclude populations where our focal *kel1*-P344T lineage won from our analysis, we still see a lower-than-expected number of *ste12* lineages winning in replays from post-*kel1* timepoints. This suggests that either the simulation parameters are not reflective of the experiment at these later time points or that there exists other low-frequency lineages containing beneficial mutations. Existence of other low-frequency lineages could also explain why we also see a lower-than-expected number of *ste12* lineages winning in Timepoint 3 populations, despite being a pre-*kel1* timepoint (Table S1). Based on sequencing of clones from the replay experiments we do see several lineages that fixed in independent populations, suggesting that there were non-*ste12*, non-focal lineages present in the original BYS2-E01 population (Figure S3). Furthermore, we expected to find the most *kel1*-P344T-containing lineages winning in replays initiated from Timepoint 4, right after the *kel1*-P344T mutation was observed and before the *ste12* lineage reached high frequency; however, no *kel1*-P344T lineages won in replays from Timepoint 4. Therefore, other mutations in the population must also be influencing the dynamics of the replay experiments.

We sequenced 50 populations from the first and second set of replay experiments and identified putative *kel1*-P344T genetic interactors as genes mutated multiple times in the *kel1*-P344T lineages. Only one of the four recurrently mutated genes, *HSL1*, was previously identified as a *KEL1* interactor using methods that assay pairwise gene deletion mutants. Interestingly, a genetic interaction between *kel1Δ* and *hsl1Δ* is absent from the Cell Map dataset (Costanzo et al. 2016), however, genetic interactions between *kel1Δ* and *hsl1Δ* have been reported elsewhere (Sharifpoor et al. 2012; Kuzmin et al. 2018). In addition, *HSL1* was identified as a recurrent target of selection in another evolution experiment initiated with a strain containing a *KEL1* mutation (Kryazhimskiy et al. 2014).

*HSL7*, the gene that we found to interact with the *kel1*-P344T mutation in Lang et al. 2013 was also identified as a recurrently mutated gene in the replay experiments. *HSL7* is a known substrate of phosphorylation by *HSL1* and association of Hsl1 and Hsl7 at the bud neck is required to advance through Swe1-mediated G2-M delay (Ma et al. 1996; Cid et al. 2001). The nonsynonymous mutations in *HSL7* that arose in the original population (A695fs) and the replay experiments (L423^*^) both result in loss of residues phosphorylated by Hsl1, while maintaining residues important for Hsl1 association (Cid et al. 2001). Two of the three mutations in *HSL1* that arose in our replay experiments (Q213K and A262P) fall within the protein kinase domain (residues 81-369), which could affect phosphorylation of *HSL7* (Finnigan et al. 2016). This may suggest the positive fitness effects of the *kel1-*P344T/*hsl7*-A695fs interaction and *kel1*-P344T/*hsl1*-A262P interaction result from similar molecular mechanisms.

Here we demonstrate the potential of experimental evolution to fish out genetic interactions to a query mutation. As a proof of principle, we use evolutionary replay experiments to identify genetic interactions with a mutant allele of *KEL1* that arose in a previous evolution experiment (Lang et al. 2013). We find that experimental evolution effectively pulls out both known and previously unknown genetic interactors, showing that additional complexity exists in genetic interaction networks that is inaccessible to knockdown or gene deletion screens. Even for the known interactors, the effect of interactions between evolved mutations is beneficial, in contrast to genetic interactions between gene deletions which typically reduce fitness (Costanzo et al. 2016), which suggests different mechanisms of actions. We show that experimental evolution, though underutilized for this purpose, is a capable of identifying growth-promoting and allele specific genetic interactions that are not easily detectable by existing methods.

## METHODS

### Evolutionary Replay Experiments

Replay experiments were initiated by resurrecting the BYS2-E01 population (described in Lang et al. 2013) at six timepoints corresponding to generations 140, 210, 270, 335, 375, and 415. These specific generations were chosen to encompass a range of timepoints before and after the *kel1*-P344T mutation was detectable. Our focal lineage was first detected by whole-genome, whole-population sequencing at Generation 140, via the initial mutation in that lineage, *iqg1*-S571N. For each timepoint, the population was resurrected by thawing the frozen stock and diluting 1:2^5^ in Yeast extract Peptone Dextrose (YPD) broth. After incubation at 30°C for 24 hours, each culture was diluted 1:2^10^ and distributed into 48 wells of a 96-well plate. The populations were then propagated in the same conditions as in the original experiment (Lang et al. 2011). Briefly, each day populations were diluted 1:2^10^ using a BiomekFX liquid handler into 128 μL of YPD plus 100 μg/mL ampicillin and 25 μg/mL tetracycline to prevent bacterial contamination. The dilution scheme equates to 10 generations of growth per day at an effective population size of ∼10^5^

### Phenotypic and Genotypic Tracking of Replay Populations

To track the dynamics of the *ste12* lineage, we measured the frequency of sterile cells in the replay experiments every 70 generations. The saturated culture was diluted 1:2^5^ into YPD plus α-factor (10 μg/mL) and incubated 6 hours at 30°C as was done previously (Lang et al. 2011). In the mating-competent ancestor, mating pheromone activates the mating pathway and induces expression of a downstream fluorescent reporter, yEVenus (Lang et al. 2009). However, cells that possess a sterile mutation do not respond to mating pheromone and do not fluoresce. Ratios of fluorescent to non-fluorescent cells was determined via flow-cytometry. Flow-cytometry data were analyzed with FlowJo version 10.5.3.

For populations where sterile frequency dropped below 0.5, two clones were isolated by streaking the population to singles on YPD agar. To identify non-sterile clones, cells were incubated with α-factor for 3 hours and visually examined for shmoos. Non-sterile clones were then grown in 5 mL YPD and a sample was stocked for further analysis. Genomic DNA was harvested using phenol-chloroform extraction and precipitated in ethanol. To identify clones derived from our focal lineage we relied on a novel *SspI* cut site (AATATT) created by a G to A transition in *IQG1* (*iqg1-*S571N). The *IGQ1* locus was amplified via PCR and incubated with *SspI* (New England Biolabs) overnight. Digested products were visualized on an agarose gel. Correct genotyping of wildtype *IQG1* and mutant *iqg1* samples was verified by submitting 10% of samples for Sanger sequencing.

### Computational Simulations

Simulations of *ste12* lineage trajectories were performed using a forward-time algorithm designed to match the conditions (including population size, dilution bottleneck, and dilution frequency) in the evolution experiment (Frenkel et al. 2014). MATLAB scripts were downloaded from https://github.com/genya/asexual-lineage-adaptation. Estimates for the distribution of beneficial fitness effects (an exponential distribution with mean = 0.85%) and beneficial mutation rate (U_b_ = 10^−4^) were used as described previously (Frenkel et al. 2014).

Simulations were performed with constant inputs for the distribution of fitness effects, beneficial mutation rate, and fitness effect of the *ste12*-Q151fs mutation, which was determined by assaying the relative fitness of *ste12* lineage clones isolated from the BYS2-E01 population (s_0_ = 3.11%) (Supplemental Dataset 2). The initial frequency of the *ste12* lineage varied (f_0_ = 0.0084, 0.0186, 0.0472, 0.2029, 0.2794, 0.4230) for each set of simulations based on estimated initial *ste12*-Q151fs frequency at each timepoint as previously reported (Lang et al. 2011). The inoculation time of the *ste12* lineage in the simulations varied (t = 0,5), as some plates were frozen 5 generations after sterile frequencies were measured in Lang et al. 2011. For each starting frequency of the *ste12* lineage, ten thousand simulations were performed. If the frequency of a simulated *ste12* lineage was greater than 0.5 at the end of the simulation, we considered it had “won”.

### Whole-Genome Sequencing

Clones were grown to saturation in 5 mL YPD and spun down to pellets and frozen at −20°C. Genomic DNA was harvested from frozen cell pellets using modified phenol-chloroform extraction and precipitated in ethanol. Total genomic DNA was used in a Nextera library preparation. The protocol used was described previously (Buskirk et al. 2017). Pooled clones were sequenced using an Illumina HiSeq 2500 sequencer by the Sequencing Core Facility at the Lewis-Sigler Institute for Integrative Genomics at Princeton. Average sequencing read depth was ∼45.

### Sequencing Analysis

Raw sequencing data were concatenated and then demultiplexed using a custom python script (barcodesplitter.py) from L. Parsons (Princeton University). Adapter sequences were trimmed using fastx_clipper from the FASTX Toolkit. Trimmed reads were aligned to a W303 reference genome (Matheson et al. 2017) using BWA v0.7.12 (Li and Durbin 2009) and mutations were called using FreeBayes v0.9.21-24-381 g840b412 (Garrison and Marth 2012). All calls were confirmed manually by viewing BAM files in IGV (Thorvaldsdóttir et al. 2013). Clones were predicted to be autodiploids if two or more mutations were called at an allele frequency of 0.5 and then manually confirmed by viewing BAM files in IGV.

### Strain Construction

We selected several putative *KEL1*-interacting mutations to reconstruct. Mutations in *HSL1, IRA2*, and *YLR001C* were chosen as they were recurrently mutated genes unique to our focal lineage. The other mutations chosen were done so based on several criteria including ease of construction (specifically, if the mutation fell within a Cas9 gRNA target sequence to avoid continual cutting by constitutively active Cas9), if there were no other candidate mutations present in that clone, and if the mutation was nonsynonymous. We also attempted to reconstruct mutations in *LTE1* (S222^*^), *BOI1* (G781D), and *CLB2* (V172fs) but were unsuccessful.

Evolved mutations were introduced into the ancestral background via CRISPR/Cas9 allele swaps as described previously (Fisher et al. 2019). Briefly, evolved mutations were reconstructed by amplifying 500bp fragments centered around the mutation of interest. These linear PCR products were transformed into the ancestor, yGIL432 (*MAT***a**, *ade2-1, CAN1, his3-11, leu2-3,112, trp1-1, URA3, bar1*Δ::*ADE2, hmlα*Δ::*LEU2, GPA1*::NatMX, *ura3*Δ::p*FUS1*-yEVenus), along with a plasmid encoding Cas9 and gRNAs targeting near the mutation site (Addgene #67638) (Laughery et al. 2015). Successful genetic reconstructions were confirmed via Sanger sequencing.

Deletion mutants were generated by integrating KanMX markers, amplified from the yeast deletion collection, into the targeted loci of yGIL432. Double deletion mutants were generated by making the KanMX deletion of *KEL1* in the *MATα* version of the ancestor and crossing it to the corresponding *MAT****a*** deletion mutants, sporulating, and selecting *MAT****a*** spores with both deletions. Integration of the KanMX cassettes at the correct loci was confirmed via Sanger sequencing.

### Competitive Fitness Assays

We measured the effect of evolved mutations on fitness using competitive fitness assays described previously (Buskirk et al. 2017; Fisher et al. 2018). Briefly, query strains are mixed 1:1 with a fluorescently-labeled version of the ancestral strain. Co-cultures are propagated in 96-well plates in the same conditions in which they evolved for 50 generations. Saturated cultures were sampled for flow-cytometry every 10 generations. Each genotype assayed was done so with at least 24 technical replicates of 2-3 biological replicates, except *kel1Δ*/*hsl1Δ, kel1Δ*/*psy2Δ*, and *kel1Δ*/*ira2Δ* which were assayed using at least 24 technical replicates of a single biological replicate each.

### Quantifying Genetic Interactions

Additive expectation of double mutant fitness effects were calculated by summing individual fitness effects of mutations and propagating uncertainty using the following formula: 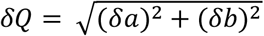, where *δa* and *δb* are the uncertainty (standard deviation) of measured fitness effects a and b. If the measured fitness effect of a double mutant differed significantly from the expected value (using Welch’s modified t-test, cutoff of p < 0.01) we considered the two mutations to interact genetically.

### Phylogenetic Analysis

The evolutionary history of our 50 sequenced clones was inferred using the Maximum Likelihood method and Tamura-Nei model (Tamura and Nei 1993). The tree with the highest log likelihood (−4514.24) was used. The tree is drawn to scale with branch lengths measured in the number of substitutions per site. Evolutionary analyses were conducted in MEGA X (Kumar et al. 2018).

## DATA AVAILIBILITY

The short-read sequencing data reported in this paper have been deposited in the NCBI BioProject database (accession no. PRJNA693025).

## ACKNOWLEDGEMENTS

We thank members of the Lang lab for comments on the manuscript. This work was supported by NIH grant 1R01GM127420 (G.I.L).

---

**Figure S1.**
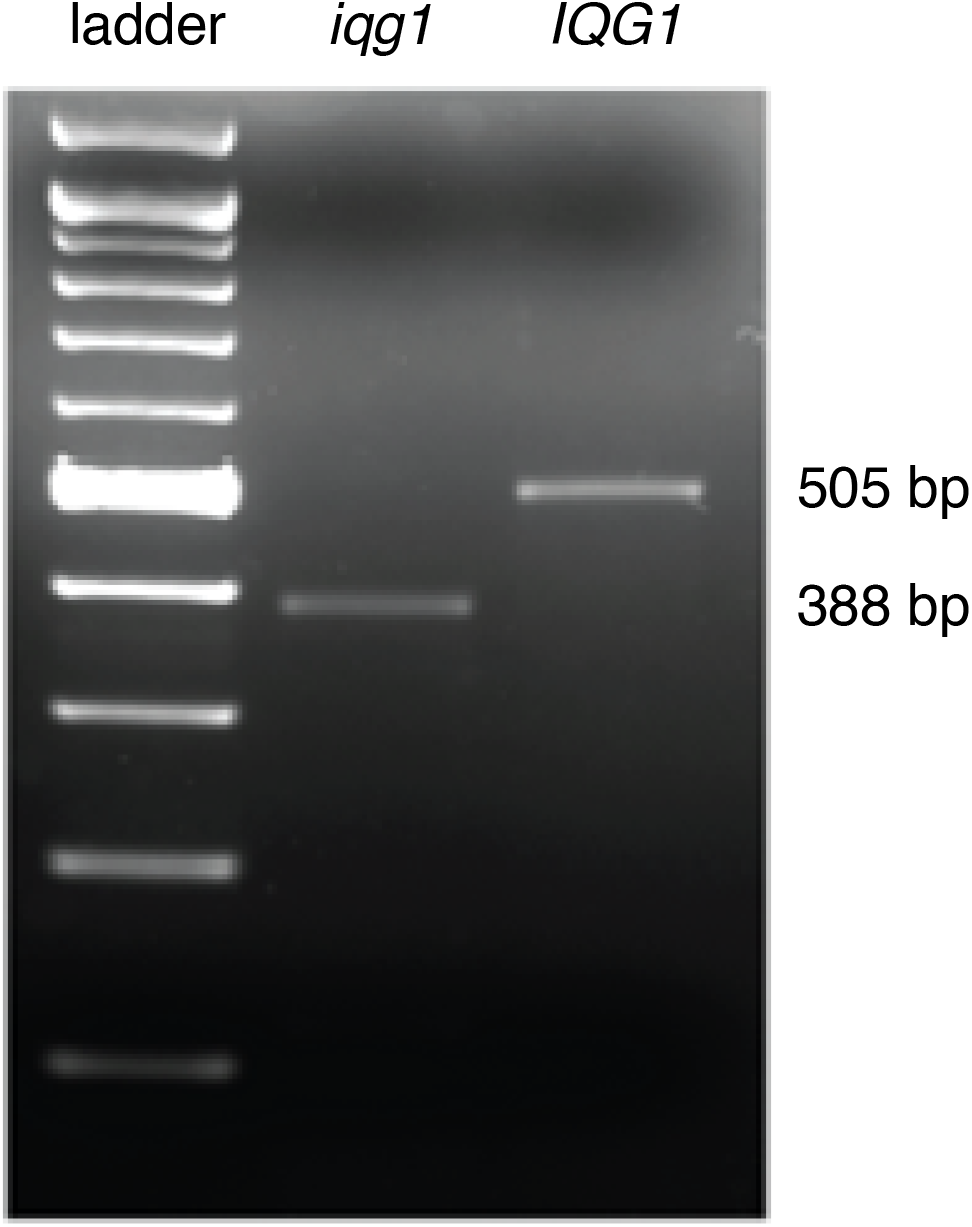
Agarose gel of SspI-digested wildtype *IQG1* and mutant *iqg1*-S571N (1712G>A) present in the focal lineage at all timepoints. Note there is a lower 117 bp band in the *iqg1* lane, but it is not easily visible.

**Figure S2.**
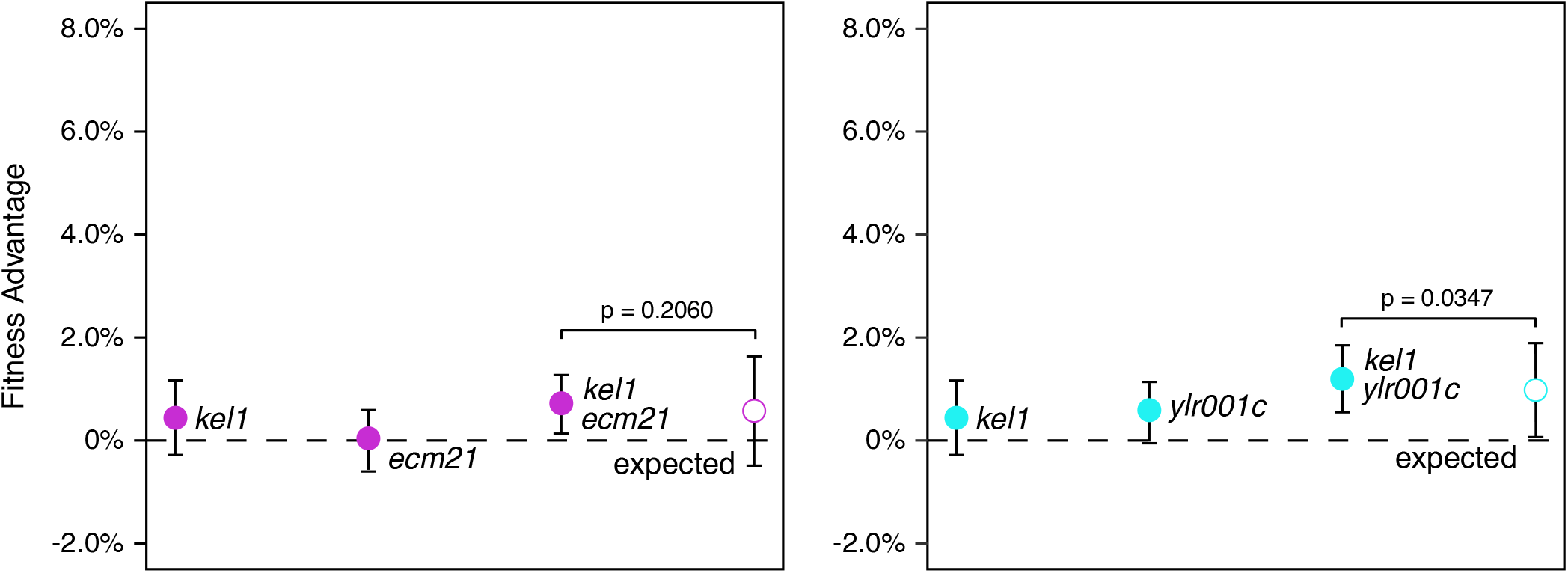
Not all mutations assayed from replay populations interacted with *kel1*-P344T. There is no detectable genetic interaction between an evolved mutation in *ecm21*-P799fs and *kel1*-P344T (left) or between *ylr001c*-R427fs and *kel1*-P344T (right).

**Figure S3.**
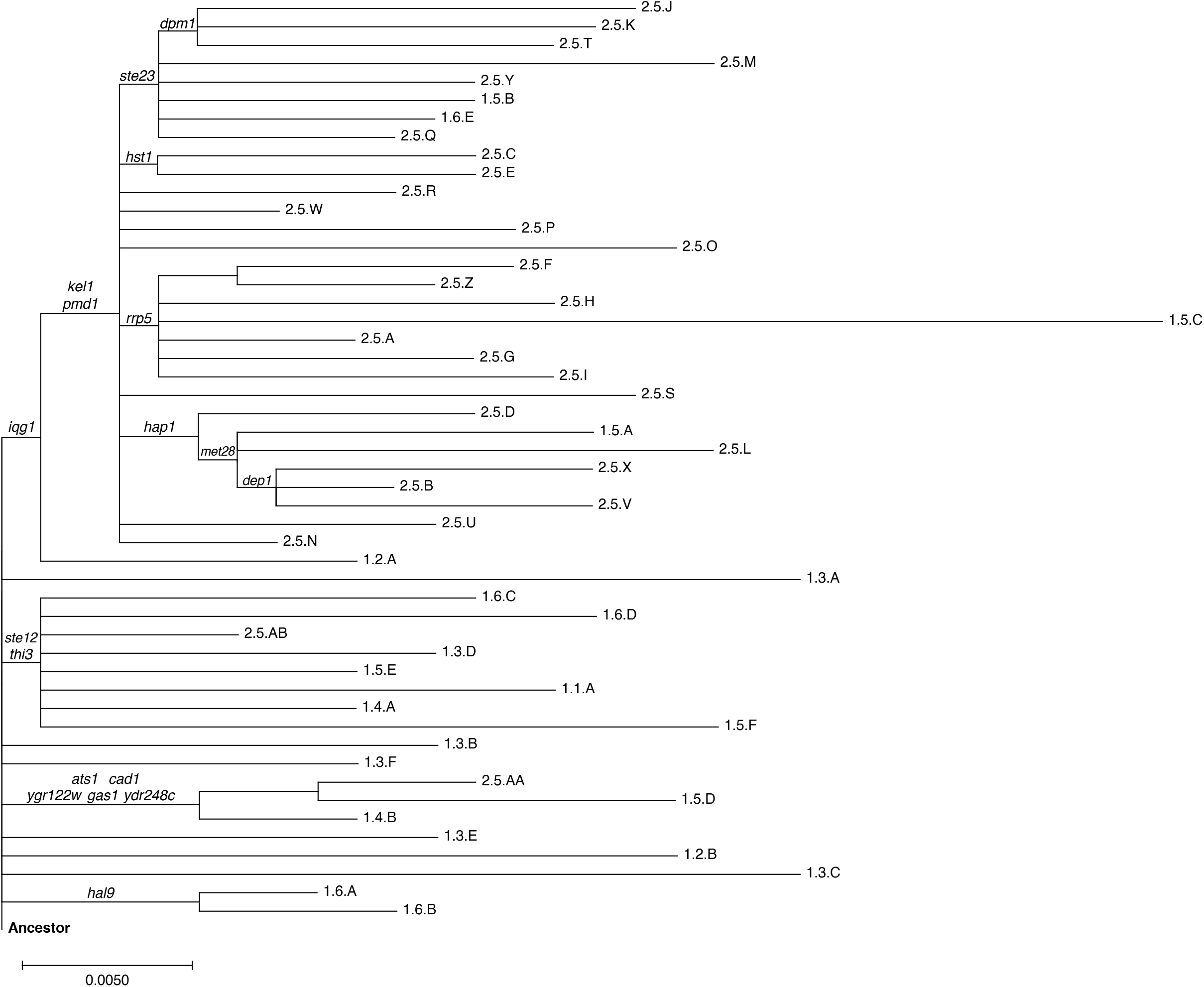
Phylogenetic tree of clones sequenced from the first round and the second round replay experiments. Common mutations (excluding intergenic mutations) are labeled above the branch on which they arose. Branch lengths are measured in the number of substitutions per site. Branch labels correspond to clone names reported in Supplemental Dataset 2; the first number corresponds to first-round (1) or second-round replay (2), the second number refers to the replay timepoint (1-6), and the letters are arbitrarily assigned to differentiate clones from the same round and timepoint but from different populations.

**Table S1.**
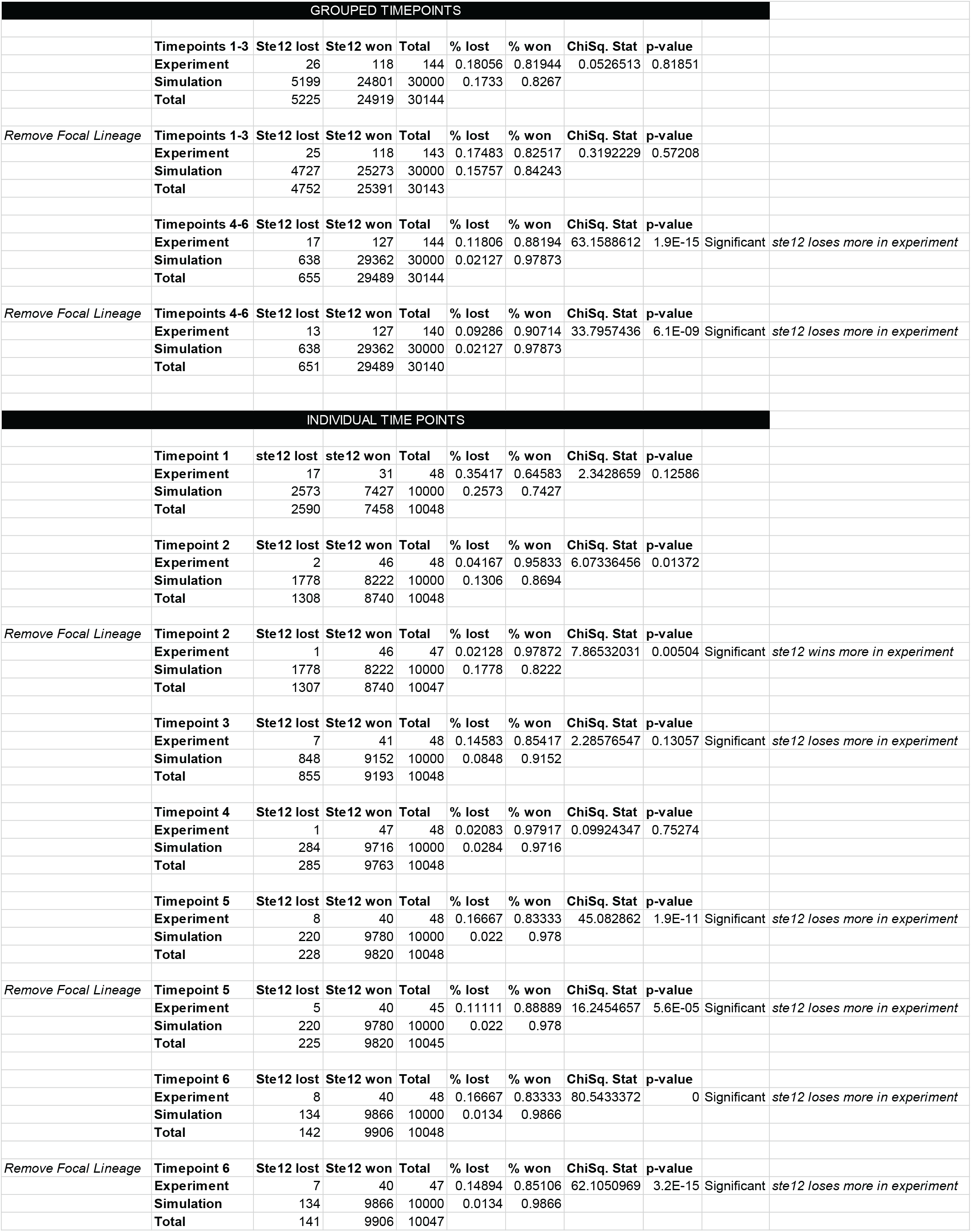

